# Phosphorylation of the novel mTOR substrate Unkempt regulates cellular morphogenesis

**DOI:** 10.1101/2022.04.08.487575

**Authors:** Pranetha Baskaran, Simeon R. Mihaylov, Elin Vinsland, Kriti Shah, Lucy Granat, Sila K. Ultanir, Andrew R. Tee, Jernej Murn, Joseph M. Bateman

## Abstract

Mechanistic target of rapamycin (mTOR) is a protein kinase that integrates multiple inputs to regulate anabolic cellular processes. mTOR complex I (mTORC1) has key functions in growth control, autophagy and metabolism. Much less is known about the signalling components that act downstream of mTORC1 that regulate cellular morphology, a vital determinant of cellular function. Here we show that the RNA-binding protein Unkempt, a key regulator of cellular morphogenesis, is a novel substrate mTORC1. We find that Unkempt phosphorylation is regulated by nutrient levels and growth factors via mTORC1. Furthermore, Unkempt physically interacts with and is directly phosphorylated by mTORC1 through binding to the regulatory-associated protein of mTOR, Raptor. Phosphorylation of Unkempt, which we find is mTORC1-dependent in cultured mammalian cell lines as well as in primary tissues, occurs largely within the highly serine-rich intrinsically disordered region of Unkempt. Importantly, mutation analysis of this region indicates that phosphorylation inhibits the ability of Unkempt to induce a bipolar morphology. Our findings reveal a novel molecular link between mTORC1 signalling and cellular morphogenesis.

## Introduction

Mechanistic target of rapamycin (mTOR) is a large serine/threonine kinase that forms the catalytic subunit of two complexes, mTOR complex 1 (mTORC1) and mTOR complex 2 (mTORC2) (Saxton & Sabatini, 2017). mTORC1 integrates signalling inputs from cellular nutrients, growth factors, energy, and stress to regulate a range of anabolic processes. Hyperactivation of mTORC1 signalling is implicated in myriad disease contexts including cancer, metabolic, immune and neurological diseases (Liu & Sabatini, 2020; Tee *et al*, 2016). mTORC2 was first shown to regulate the actin cytoskeleton via the AGC kinase PKCα and was subsequently shown to have additional roles in cell proliferation and survival (Jacinto *et al*, 2004; Sarbassov *et al*, 2004; Saxton & Sabatini, 2017).

mTOR regulates the phosphorylation of several hundred proteins, dependent on the cell type (Battaglioni *et al*, 2022; Demirkan *et al*, 2011; Hsu *et al*, 2011; Robitaille *et al*, 2013; Yu *et al*, 2011). Relatively few direct mTOR substrates have however been characterised in detail. Well characterised mTORC1 substrates include S6K, 4E-BP1 and the adaptor protein Grb10, all regulators of cell growth and the protein kinase ULK1, which regulates autophagosome biogenesis (Ben-Sahra *et al*, 2013; Egan *et al*, 2011; Hsu *et al*., 2011; Saxton & Sabatini, 2017; Yu *et al*., 2011). mTORC2 phosphorylates AKT and SGK1 to control cell proliferation and survival (García-Martínez & Alessi, 2008; Sarbassov *et al*., 2004; Sarbassov *et al*, 2006). mTORC2 also phosphorylates multiple PKC isoforms, including PKCα, PKBβII and PKCζ, to regulate actin cytoskeletal dynamics (Baffi *et al*, 2021; Ikenoue *et al*, 2008; Li & Gao, 2014).

In contrast to mTORC2, the role of mTORC1 in cellular morphogenesis is much less well understood. We previously showed that the zinc finger/RING domain protein Unkempt acts genetically downstream of mTOR in the developing retina and central nervous system in *Drosophila* (Avet-Rochex *et al*, 2014; Bateman, 2015; Bateman & McNeill, 2004; Maierbrugger *et al*, 2020). Mammalian Unkempt is a key regulator of cell morphogenesis and, through its zinc finger domain, binds mRNAs involved in control of protein synthesis and cell shape to affect their translation and to induce the establishment of the early neuronal morphology (Murn *et al*, 2016; Murn *et al*, 2015).

Here we show that mammalian Unkempt is abundantly phosphorylated and that phosphorylation is acutely sensitive to mTORC1 activity. Unkempt physically interacts with mTORC1 by binding to regulatory-associated protein of mTOR (Raptor), which mediates direct phosphorylation by mTOR at multiple serine/threonine residues in the highly serine-rich intrinsically disordered region (IDR). Using mice with conditional knockout of *Tsc1* to activate mTORC1 in the nervous system, we find that Unkempt phosphorylation is mTORC1 dependent *in vivo*. Finally, we show that phosphorylation in the serine rich region suppresses the capacity of Unkempt to induce a bipolar morphology, a critical cellular activity of Unkempt.

## Results and Discussion

### mTORC1 activity regulates Unkempt phosphorylation

Unkempt is a highly conserved RNA-binding protein that plays critical roles in controlling cell proliferation and differentiation in *Drosophila* and cellular morphogenesis in mammals (Avet-Rochex *et al*., 2014; Li *et al*, 2019; Maierbrugger *et al*., 2020; Murn *et al*., 2015). How the activity of Unkempt might be regulated remains unknown. Recent studies have suggested that Unkempt acts genetically downstream of mTORC1 (Avet-Rochex *et al*., 2014; Li *et al*., 2019; Maierbrugger *et al*., 2020), but it is unclear whether mTORC1 signalling might be linked to the function of Unkempt. Mammalian Unkempt is either not expressed or expressed at very low levels in non-neural cell lines but is abundant in cells of neuronal lineages (Murn *et al*., 2015). We therefore used human neuroblastoma SH-SY5Y cells to determine whether Unkempt is regulated in an mTORC1-dependent manner.

In serum-starved neuroblastoma SH-SY5Y cells we found that stimulation of the mTORC1 pathway with insulin or serum re-addition caused a mobility shift of Unkempt to a higher resolved band, whereas direct inhibition of mTORC1 kinase activity with rapamycin prevented this shift (Figure 1A). A strict correlation between inhibition of mTORC1, as judged by phosphorylation of ribosomal protein S6 (rpS6), and decreased mobility of Unkempt was observed over a wide range of rapamycin concentrations (Supplemental Figure 1A). Moreover, a phospho-specific antibody against Unkempt (see details below and Figure 3D) detected reduced levels of phosphorylated Unkempt in rapamycin treated cells (Supplemental Figure 1A). Rapamycin treatment also caused increased phosphorylation of AKT at Ser473 as a result of the established feedback loop between S6K and the insulin receptor substrate (IRS) (Harrington *et al*, 2004; Shah *et al*, 2004) (Supplemental Figure 1A). Unkempt showed a similar mTORC1-dependent mobility shift in response to rapamycin in mouse N2a neuroblastoma cells (Figure 1B) and in mouse E14.5 primary cortical cells (Figure 1C). After insulin stimulation, lambda protein phosphatase treatment of the SH-SY5Y cell lysate converted a higher resolving Unkempt band to a lower resolving species, below that in unstimulated cells (Figure 1D). Together, these data suggest that Unkempt is phosphorylated in an mTORC1-dependent manner and exhibits a basal level of phosphorylation in unstimulated cells.

**Figure 1.**
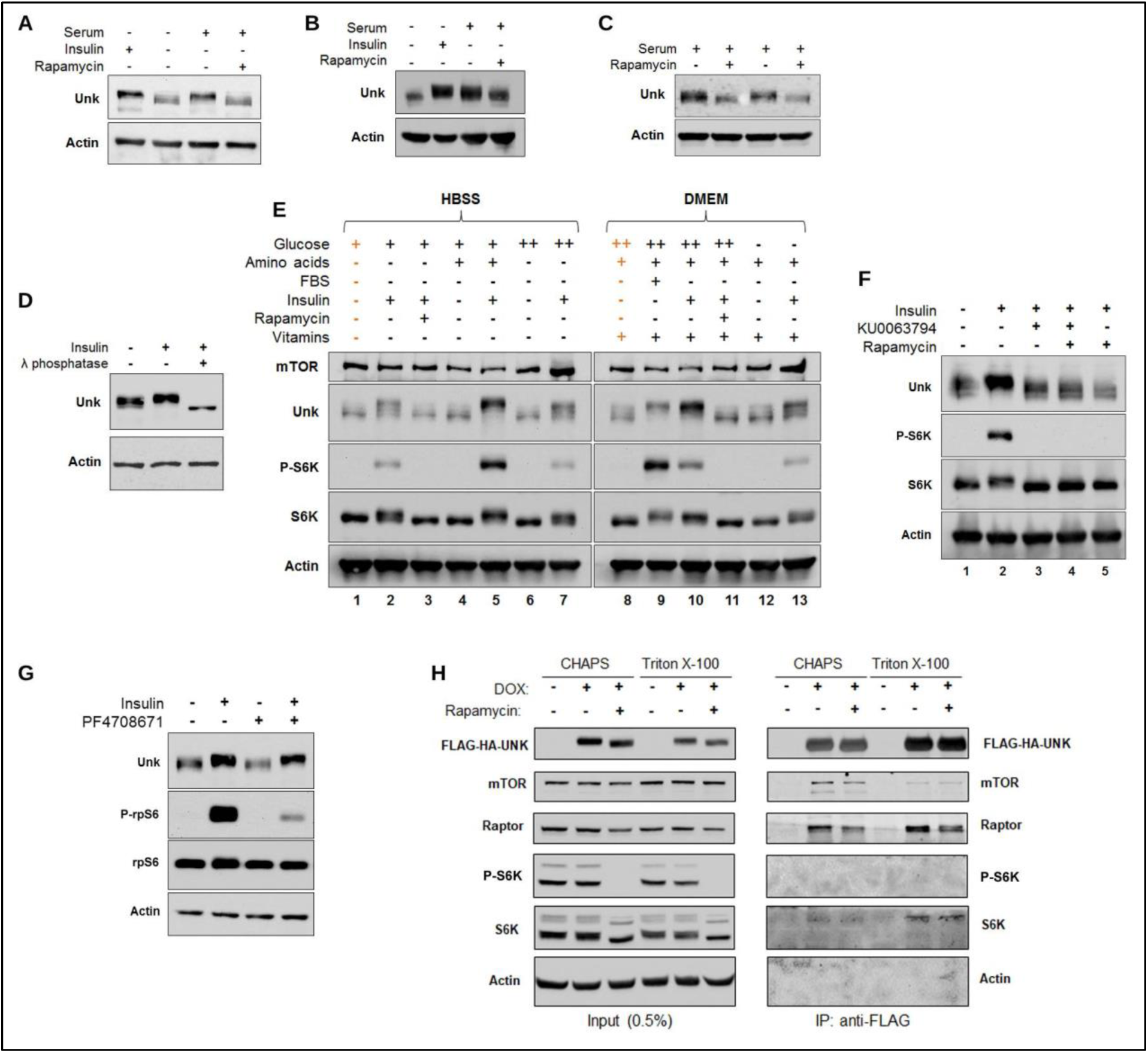
Phosphorylation of Unkempt is mTORC1 activity dependent and physically Unkempt interacts with Raptor. (A) Western blot showing serum starvation followed by insulin stimulation of SH-SY5Y cells increases, while rapamycin treatment decreases the electrophoretic mobility of Unkempt. (B, C) Rapamycin treatment decreases the electrophoretic mobility of Unkempt in mouse N2a cells and mouse E14.5 primary cortical cells. (D) Treatment of SH-SY5Y cell lysate with lambda protein phosphatase abrogates the electrophoretic mobility shift in Unkempt. (E) Unkempt phosphorylation is highly sensitive to nutrient levels. SH-SY5Y cells were grown in minimal media lacking vitamins and amino acids (HBBS) or DMEM. Additional nutrients were added as indicated. Orange columns indicate control for that panel. (F) Treatment of SH-SY5Y cells with KU0063794 alone, rapamycin alone, or both KU0063794 and rapamycin all prevent the increase in Unkempt phosphorylation caused by insulin stimulation. (G) Inhibition of S6K with PF4708671 does not prevent the phosphorylation of Unkempt. (H) Unkempt physically interacts with Raptor and mTOR. Mock purified extract from un-induced HeLa S3 cells (DOX -) was used as a negative control.

mTORC1 can be regulated by multiple signalling inputs, including insulin, amino acids and ATP levels (Saxton & Sabatini, 2017). We found that addition of amino acids alone in the media did not suffice to activate mTORC1, as judged by phosphorylation of S6K, or increase Unkempt phosphorylation (Figure 1E, compare lanes 1 and 4). However, the presence of amino acids amplified the effect of insulin stimulation, as seen both by increased mTORC1 activity and enhanced phosphorylation of Unkempt (Figure 1E, compare lanes 2 and 5). Similarly, increased glucose concentration, while having little effect on its own (Figure 1E, compare lanes 8 and 12), potentiated mTORC1 activity and the phosphorylation of Unkempt primed by insulin stimulation in the presence of amino acids (Figure 1E, compare lanes 10 and 13). The effects of amino acids and glucose were mTORC1-dependent, as rapamycin prevented the phosphorylation of Unkempt in the presence of amino acids, glucose, and insulin together (Figure 1E, compare lanes 10 and 11). Unkempt phosphorylation is therefore acutely sensitive to nutrients and growth factors that regulate mTORC1.

mTOR is a component of both mTORC1 and mTORC2. Prolonged rapamycin treatment can inhibit mTORC2 (Sarbassov *et al*., 2006) but short-term (one hour) rapamycin treatment of SH-SY5Y cells did not inhibit mTORC2, as judged by AKT phosphorylation at Ser473 (Supplemental Figure 1A). Moreover, treatment of SH-SY5Y cells with a dual mTORC1/mTORC2 inhibitor, KU0063794 (Figure 1F, lane 3), or rapamycin and KU0063794 together (Figure 1F, lane 4), produced a similar reduction in phosphorylation of Unkempt as treatment with rapamycin alone (Figure 1F, lane 5), further evidence that it is the activity of mTORC1 but not mTORC2 that regulates Unkempt phosphorylation. Furthermore, treatment of SH-SY5Y cells with a range of concentrations of the highly selective mTORC1 inhibitor DL001 (Schreiber *et al*, 2019), which does not inhibit mTORC2-dependent AKT phosphorylation, decreased the mobility of Unkempt and reduced the recognition by an Unkempt phospho-specific antibody (Supplemental Figure 1B). We also asked whether Unkempt phosphorylation might be regulated at the level of S6K, however, treatment of SH-SY5Y cells with the S6K inhibitor PF4708671 had no noticeable effect (Figure 1G). Together, these data confirm that Unkempt phosphorylation is mTORC1-dependent.

### Unkempt physically interacts with mTORC1

Unkempt has been previously shown to physically interact with Raptor (Li *et al*., 2019; Vinsland *et al*, 2021). To further test whether Unkempt physically interacts with mTORC1 we used HeLa S3 cells with a doxycycline (DOX)-inducible FLAG-HA-Unkempt to immunoprecipitate Unkempt (Murn *et al*., 2015). Rapamycin treatment showed that phosphorylation of FLAG-HA-Unkempt is mTORC1-dependent (Figure 1H). Endogenous Raptor and mTOR co-immunoprecipitated with FLAG-HA-Unkempt (Figure 1H). Unkempt therefore physically interacts with mTORC1.

We also noticed that the co-immunoprecipitation of mTOR in this experiment was sensitive to the type of detergent, whereas pulldown of Raptor was not (Figure 1H). This is in line with a known property of the zwitterionic detergent CHAPS to preserve and the non-ionic detergent Triton X-100 to disrupt the interaction between Raptor and mTOR (Kim *et al*, 2002), consistent with Unkempt interacting with mTOR indirectly via Raptor. These co-immunoprecipitation data further validate the previously observed physical interaction of Unkempt with Raptor (Li *et al*., 2019; Vinsland *et al*., 2021) and show that Unkempt physically interacts with a functional mTORC1.

### mTORC1 phosphorylates Unkempt at multiple sites in its intrinsically disordered region

Using mass spectrometry (MS) of FLAG-HA-Unkempt immunoprecipitated from cells treated or not with rapamycin we identified 31 phosphorylated serines and five phosphorylated threonines grouped in three clusters within a predicted intrinsically disordered region (IDR) of Unkempt (Figure 2A, B, Supplemental Figures S2, S3). About 24% of the C-terminal portion of the IDR between amino acids 467 and 640 consists of serine residues (41 residues) and harbours the majority of phosphorylation sites (phosphosites), including all of the phosphosites that were never detected in rapamycin-treated cells (Figure 2A, blue residues, Supplemental Figures S2, S3). To determine if any of the phosphosites might be direct substrates of mTORC1 we used an *in vitro* kinase assay with purified reconstituted mTORC1 and GTP-bound Rheb. We found that purified Unkempt was indeed directly phosphorylated by mTORC1/Rheb-GTP (Figure 2C). MS identified five serine residues in Unkempt that were *in vitro* phosphorylated in the control condition but not in the rapamycin/FKBP12 treated condition (addition of the rapamycin binding partner FKBP12 is necessary for mTORC1 inhibition *in vitro* (Dunlop *et al*, 2009); Figure 3A, red asterisks; Supplemental Figures S2, S3). Four of these serines have a proline or leucine at the +1 position and so adhere to the predicted mTORC1 phosphorylation motif consensus sequence (Hsu *et al*., 2011; Robitaille *et al*., 2013), while two (S608 and S611) are also mTORC1-dependent in rapamycin-treated cells (Figure 2A, Supplemental Figures S2, S3).

**Figure 2.**
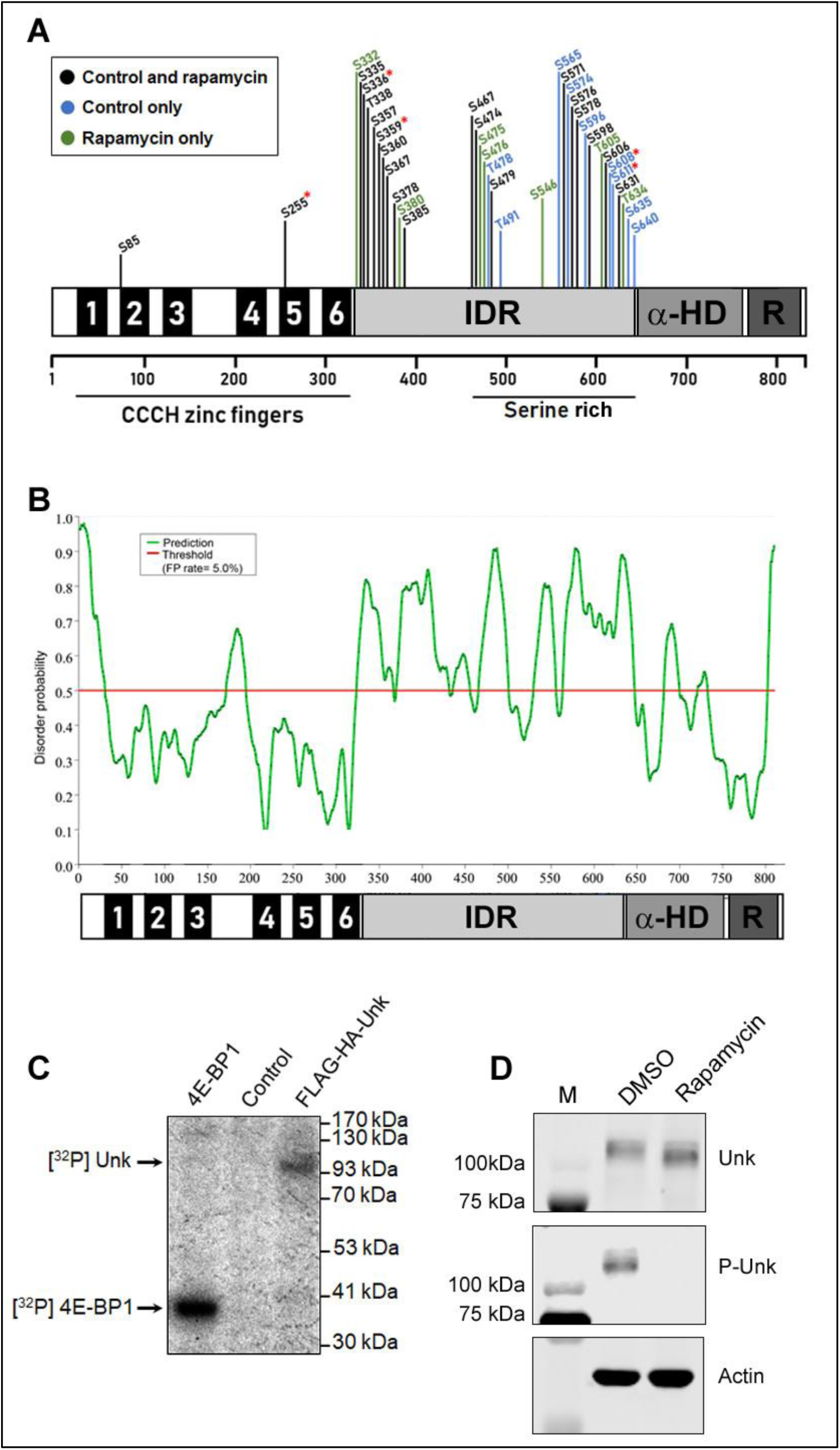
Unkempt is a direct substrate of mTORC1 and is phosphorylated in the IDR by mTORC1. (A) A schematic of the phosphorylated residues identified in Unkempt by LC-MS/MS. Phosphorylated residues identified in both control and rapamycin treated cells shown in black; phosphorylated residues identified only in control cells shown in blue; phosphorylated residues identified only in rapamycin treated cells shown in green. Phosphorylated residues identified only in the control *in vitro* kinase assay condition and not in the rapamycin/FKBP12 condition are highlighted with a red asterisk. See Supplemental Figures S2 and S3 for details. IDR: intrinsically disordered region, α-HD: alpha-helical domain, R: ring domain. (B) Prediction disorder probability plot of the primary sequence of mouse Unkempt. The serine-rich region (amino acids 467 to 640) is highly disordered. False positive rate of 5%. (C) FLAG-HA-Unkempt purified from HeLa S3 cells is phosphorylated *in vitro* by reconstituted mTORC1/Rheb-GTP. Recombinant 4E-BP1 was used as a positive control. Mock purified extract from un-induced HeLa S3 cells was used as a negative control. (D) A phospho-specific antibody raised against phospho-S606/ phospho-S611 in Unkempt does not recognise Unkempt from rapamycin treated SH-SY5Y cells. M: molecular weight marker.

**Figure 3.**
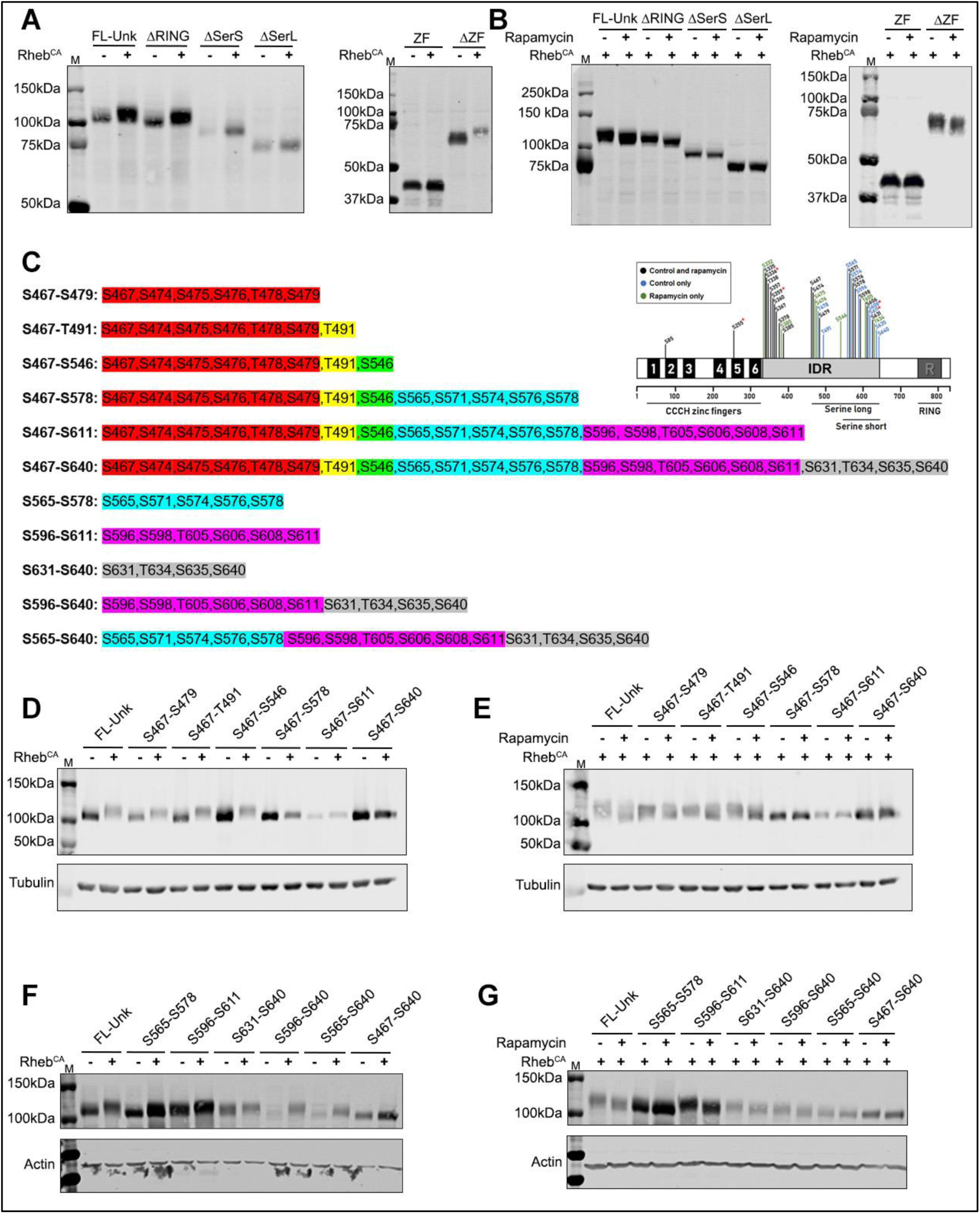
Phosphorylation within the serine-rich region of the IDR is necessary for mTORC1 dependent phosphorylation of Unkempt. (A, B) Full length V5-Unkempt (FL-Unk), or deletion mutants lacking the zinc finger domain (ΔZF) or the RING domain (ΔRING) are phosphorylated in HEK293 cells expressing constitutively active Rheb (Rheb^CA^) (A) and dephosphorylated in these cells treated with rapamycin (B). Phosphorylation is absent with zing finger domain alone (ZF) and completely or partially absent with deletion mutants lacking whole serine-rich region (ΔSerL) or part of the serine-rich region (ΔSerS) (A, B). (C) A schematic of the phosphorylated residues identified in Unkempt by LC-MS/MS. Phosphorylated residues identified in both control and rapamycin treated cells shown in black; phosphorylated residues identified only in control cells shown in blue; phosphorylated residues identified only in rapamycin treated cells shown in green. Phosphorylated residues identified only in the control *in vitro* kinase assay condition and not in the rapamycin/FKBP12 condition are highlighted with a red asterisk. See Supplemental Figures S1 and S2 for details. (D, E) Full length V5-Unkempt (FL-Unk) or FL-Unk containing alanine mutations in phosphorylated serine and threonine residues between S467-S546 expressed in HEK293 cells are phosphorylated by constitutively active Rheb (Rheb^CA^) (D) and dephosphorylated by rapamycin (E). Additional mutation of phosphorylated residues from S467-S578, S467-S611 and S467-S640 partially or completely prevent phosphorylation by constitutively active Rheb (D) and dephosphorylation by rapamycin (E). (F, G) Alanine mutants of Unkempt in phosphorylated serine and threonine residues between S565-S640 are phosphorylated by constitutively active Rheb (Rheb^CA^) (F) and dephosphorylated by rapamycin (G). M: molecular weight marker.

Unkempt phosphorylation at S611 was reproducibly rapamycin dependent in cells (in three biological replicates) and in the *in vitro* kinase assay (Supplemental Figure S3). S611 also has a leucine residue at the +1 position and hydrophobic residues at -1 (glycine) and -4 (alanine) and so adheres to the mTORC1 phosphorylation motif consensus sequence (Hsu *et al*., 2011; Robitaille *et al*., 2013) (Supplemental Figure S2). We generated a phospho-specific antibody against a peptide containing rapamycin sensitive phospho-S611 and nearby rapamycin insensitive phospho-S606, which was also reproducibly identified in cells by MS (Supplemental Figure S3). This antibody recognised Unkempt in serum stimulated SH-SY5Y cells but not in cells that were treated with rapamycin (Figure 2D, see also Supplemental Figure S1A). Taken together, these data provide evidence that Unkempt is an mTORC1 substrate that is phosphorylated at multiple residues, located mostly in the serine-rich region of the IDR.

To independently define the region of Unkempt that is targeted by mTORC1-dependent phosphorylation we performed deletion analysis of Unkempt transiently expressed in HEK293 cells. Co-expression of constitutively active Rheb (Rheb^CA^), which directly hyperactivates mTORC1 (Hartman *et al*, 2013), caused phosphorylation of full-length Unkempt (FL-Unk) in the absence but not in the presence of rapamycin (Figure 3A, B). Unkempt deletion mutants lacking either the zinc finger domain (ΔZF) or the RING domain (ΔRING) retained mTORC1-dependent phosphorylation potential, while a mutant containing just the zinc finger domain alone (ZF) was not phosphorylated by mTORC1 hyperactivation (Figure 3A, B). Notably, removal of the whole serine-rich region (ΔSerL), or of a shorter region between amino acids 539 and 640 (ΔSerS) almost completely prevented phosphorylation (Figure 3A, B). This deletion analysis confirms that the serine-rich region in Unkempt contains the majority of the mTORC1-dependent phosphosites.

MS analysis showed that specific serine and threonine residues in the IDR of Unkempt are targets of mTORC1-dependent phosphorylation (Figure 2A). To test these data further we mutated clusters of MS-identified phosphorylated serine and threonine residues in the IDR to alanine (Figure 3C). Mutation of phosphosites from S467 to S546 to alanine had no obvious effect on mTORC1-dependent phosphorylation of Unkempt (Figure 3D, E). However, additional mutation of phosphosites between S565-S578, S596-S611 and S631-S640 to alanine either strongly reduced or completely prevented mTORC1-dependent phosphorylation of Unkempt (Figure 3D, E). Based on this, we then tested mutants that only affect phosphosites located exclusively within the serine-rich region of with the IDR (Figure 3C). Interestingly, mutations of groups of these phosphosites (S565-S578, S596-S611 or S631-S640), or their combinations (S596-S640 or S565-S640), reduced but did not completely prevent the overall mTORC1-dependent phosphorylation of Unkempt (Figure 3C, F, G). These mutagenesis data validate the MS analyses and point to the serine-rich region of the IDR as the hotspot of mTORC1-dependent phosphorylation of Unkempt.

### Unkempt is phosphorylated by mTORC1 in vivo

In mice, Unkempt protein is expressed in the developing nervous system from at least as early as E10 (Murn *et al*., 2015). To analyse the phosphorylation status of Unkempt by mTORC1 *in vivo* we utilised *Nestin-Cre;Tsc1*^*fl/fl*^ mice, in which *Nestin* promoter-driven Cre recombinase is expressed in neural progenitors beginning at E10.5 causing loss of TSC1 expression throughout the developing brain (Anderl *et al*, 2011). Consequently, *Nestin-Cre;Tsc1*^*fl/fl*^ mice exhibit activated mTORC1 signalling in the brain, macrocephaly and die at P0, likely due to malnutrition, hypoglycemia and hypothermia (Anderl *et al*., 2011). The activated mTORC1 signalling and perinatal death of these mice can be suppressed by a single dose of rapamycin between E15-E17 (Anderl *et al*., 2011). We confirmed hyperactivation of mTORC1 via increased phosphorylation of rpS6 in the brain of E16.5 *Nestin-Cre;Tsc1*^*fl/fl*^ mice embryos (Figure 4A, B). Although we did not detect a change in the electrophoretic mobility of Unkempt protein, using the phospho-specific antibody we observed a dramatic increase in Unkempt phosphorylation in brain tissue from *Nestin-Cre;Tsc1*^*fl/fl*^ embryos (Figure 4A, C).

**Figure 4.**
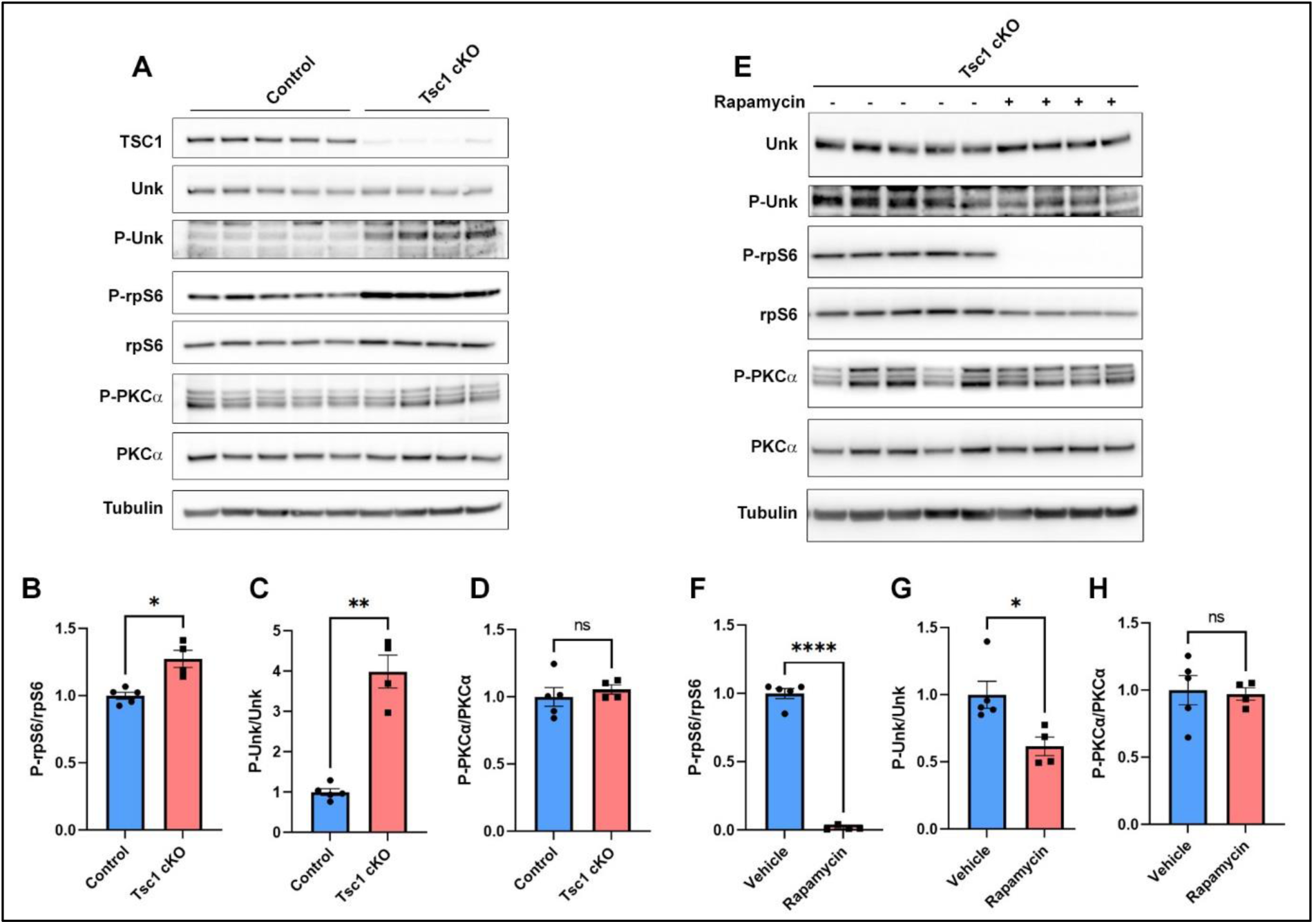
Unkempt is phosphorylated by mTORC1 during neurodevelopment. (A) Western blot analysis of E16.5 brain lysate from control (*Nestin-Cre;Tsc1*^*fl/+*^) and Tsc1 cKO (*Nestin-Cre;Tsc1*^*fl/fl*^) mice. (B-D) Quantification of phospho-rpS6 (B), phospho-Unkempt (C) and phospho-PKCα (D) expression. Control n=5, Tsc1 cKO n=4. (E) Western blot analysis of E16.5 brain lysate from Tsc1 cKO (*Nestin-Cre;Tsc1*^*fl/fl*^) mice treated with vehicle or rapamycin for 24 hours. (F-H) Quantification of phospho-rpS6 (F), phospho-Unkempt (G) and phospho-PKCα (H) expression. Vehicle n=5, rapamycin treated n=4. Data are represented as mean ± SEM. Student’s t test, ns not significant, *p<0.05, **p<0.01, ****p < 0.0001.

To test whether inhibition of mTORC1 affects Unkempt phosphorylation during nervous system development, mice pregnant with *Nestin-Cre;Tsc1*^*fl/fl*^ embryos were injected with rapamycin or vehicle at E15.5. Twenty four hours after rapamycin injection, rpS6 phosphorylation was dramatically reduced in the embryonic brain (Figure 4E, F). Rapamycin treatment also significantly decreased phosphorylation of Unkempt (Figure 4E, G). To determine whether prolonged rapamycin treatment affected mTORC2 signalling (Sarbassov *et al*., 2006), we analysed PKCα phosphorylation (Huang *et al*, 2009). PKCα phosphorylation was unnaffected in the brain of *Nestin-Cre;Tsc1*^*fl/fl*^ embryos compared to controls (Figure 4A, D) and did not change with twenty-four-hour rapamycin treatment (Figure 4E, H), indicating that mTORC2 signalling was unaffected. These data show that Unkempt phosphorylation is mTORC1 dependent in the developing brain.

### Phosphorylation of Unkempt affects its capacity to remodel cellular morphology

In addition to being required for the early morphology of neurons, Unkempt also possesses a remarkable capacity to establish a similar, early neuronal-like bipolar morphology in cells of non-neuronal origin, indicating that Unkempt regulates a specific program of cell morphogenesis (Murn *et al*., 2016; Murn *et al*., 2015). To examine whether phosphorylation affects the morphogenetic activity of Unkempt, we inducibly expressed in HeLa cells GFP either alone or together with wild-type or phosphosite-mutant Unkempt and monitored the effects on cell shape. With clusters of phosphosites turned to either alanines or aspartates to prevent or mimic phosphorylation, respectively, we focused on the serine-rich region of IDR, in particular its C-terminal region (S596-S640), as the hotspot for mTORC1-dependent phosphorylation of Unkempt (Figure 4C-G, Figure 5A). Strikingly, whereas alanine mutations largely preserved the morphogenetic activity of Unkempt, aspartate mutations severely compromised or abolished cell polarization by Unkempt (Figure 5B-D). All mutants showed similar gross RNA-binding to wild-type Unkempt (Supplemental Figure S4), suggesting that the morphogenetic effects of these mutations are unlikely to be due to differences in RNA binding, a key requirement for Unkempt-induced cell morphogenesis (Murn *et al*., 2016; Murn *et al*., 2015). Together, these data point to a critical role of phosphorylation in regulating the activity of Unkempt in cellular morphogenesis (Figure 5E).

**Figure 5.**
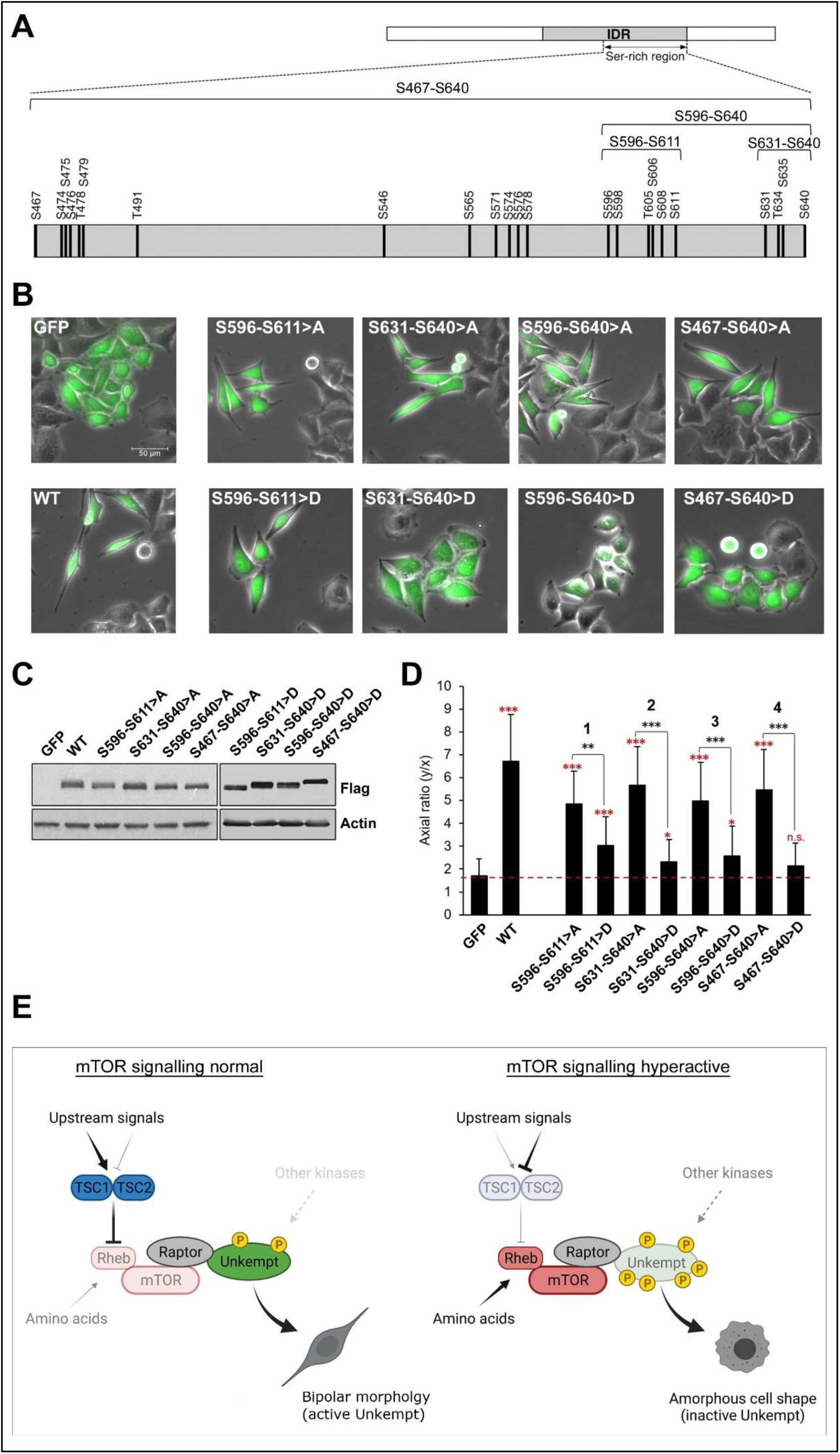
Phosphosite mutations in the serine-rich region compromise the morphogenetic activity of Unkempt. (A) Map of Unkempt protein (top) with the locations of all detected phosphosites within the serine-rich region of the IDR shown magnified on the bottom, drawn to scale. (B) GFP only-inducible or GFP and wild-type (WT) or phosphosite alanine (>A) or aspartate (>D) mutant Unkempt-inducible HeLa cells at 48 hours of treatment with doxycycline. Scale bar, 50 μm. (C) Detection by immunoblotting of WT or the indicated mutant Unkempt proteins in total cell lysates of inducible HeLa cells at 48 hours of treatment with doxycycline. (D) The morphologies of the HeLa cell cultures indicated in (B) were quantified by calculating their axial ratios (y/x), as described previously (see Materials and methods (Murn *et al*., 2015)). The results are compared with the GFP control (red asterisks) morphology, or mutant Unkempt-expressing cells are compared for each phosphosite region indicated in (A), (black asterisks). Red dashed line denotes the average axial ratio of the GFP control cells. Error bars indicate SD (n = 50). (*) 4.3E-5 < *P* < 5.2E-5, (**) *P* = 3.9E-6, (***) *P* < 1.3E-10 (pertains to black and red asterisks), Student’s *t*-test. (E) A model for the regulation of cellular morphogenesis by Unkempt acting downstream of mTORC1.

We have shown that Unkempt phosphorylation is exquisitely sensitive to mTORC1 activity and is modulated by a range of inputs including glucose, amino acids and insulin signalling. mTORC1 regulates the phosphorylation of hundreds of different proteins (Battaglioni *et al*., 2022; Hsu *et al*., 2011; Robitaille *et al*., 2013; Yu *et al*., 2011) with roles in growth control, autophagy, nucleotide and lipid metabolism but the control of cellular morphogenesis by mTORC1 is less well characterised. The microtubule associated proteins CLIP-170 and Tau are known to be direct substrates of mTORC1 but do not have the Unkempt’s profound ability to reorganise cell morphology (Choi *et al*, 2002; Tang *et al*, 2013).

Raptor physically interacts with Unkempt, enabling mTOR to phosphorylate multiple residues within the serine-rich region of the Unkempt IDR. Unkempt phosphorylation is complex and includes mTORC1-dependent and independent serine and threonine residues, suggesting the involvement of additional kinases. The number and proximity of phosphosites in the IDR also implies potential flexibility in the regulation of Unkempt phosphorylation. *In vivo* phosphorylation of Unkempt may depend on the context, both spatially and temporally. Thus, through phosphorylation, the IDR of Unkempt might serve as a rheostat that integrates inputs from multiple signalling pathways to fine-tune the activity of Unkempt (Figure 5E). This might not be limited to the control of the cellular morphogenesis but could extend to other known roles of Unkempt in neuronal differentiation and cell proliferation (Avet-Rochex *et al*., 2014; Li *et al*., 2019; Lores *et al*, 2010; Maierbrugger *et al*., 2020; Mohler *et al*, 1992; Vinsland *et al*., 2021).

It remains unclear how phosphorylation might regulate the activity of Unkempt at the molecular level. Thus far, the most extensively documented molecular property of Unkempt is its capacity to bind mRNA, which occurs in an RNA sequence-specific manner and is mediated by a tandem array of six highly conserved CCCH-type zinc fingers located at the N-terminus (Murn *et al*., 2016; Murn *et al*., 2015). Through RNA binding, Unkempt is thought to primarily regulate the translation of several hundred of its target mRNAs involved in processes including control of protein synthesis and cell morphogenesis, although the mechanistic basis for translational control and potential other modes of post-transcriptional processing by Unkempt remains unknown. Given the physical separation of the zinc finger domain from the heavily phosphorylated serine-rich region in the IDR, but also because the zinc finger domain alone suffices for high-affinity RNA binding, it seems unlikely that phosphorylation in the serine-rich region would affect RNA binding by Unkempt (Murn *et al*., 2016; Murn *et al*., 2015). Indeed, our mutagenesis analysis coupled with CLIP data suggest that phosphorylation exerts no overt effect on Unkempt-RNA interactions. Instead, we propose that phosphorylation by mTORC1 acts to modulate protein-protein interactions (PPIs) between the IDR and partner proteins, in keeping with the known propensity of IDRs to act as protein rheostats by forming transient PPIs regulated by phosphorylation (Wright & Dyson, 2015). Phosphorylation-sensitive PPIs might entail interactions of Unkempt with a large number of cytosolic RNA granule components and other effectors of post-transcriptional processing identified via unbiased screens (Go *et al*, 2021; Havugimana *et al*, 2012; Youn *et al*, 2018). Such a mechanism could provide a novel route for the mTOR pathway to control morphogenesis and other cellular processes, beyond those regulated by 4E-BP/eIF4E and S6K (Thoreen *et al*, 2012). Overall, this study paves the way for investigation of the mechanistic principles by which Unkempt and its phosphorylation by mTORC1, and potentially other kinases, regulate cellular morphogenesis.

## Acknowledgements

We are grateful to Angelique Bordey for the Rheb^CA^ plasmid, Stelios Tzannis and Aeovian Pharmaceuticals for providing DL001. This work was funded by a Health Schools PhD studentship from King’s College London to EV. The Wellcome Trust (200668/Z/16/Z) and the USA Department of Defence (W81XWH-17-1-0082) to JB. Health and Care Research Wales (the Wales Cancer Research Centre) and the Tuberous Sclerosis Association supported AT. SKU and SRM received funding from the Francis Crick Institute, which receives its core funding from Cancer Research UK (FC001201), the UK Medical Research Council (FC001201), and the Wellcome Trust (FC001201). The National Institute of General Medical Sciences grant GM144693 and the University of California Research Initiatives grant CRN-20-637684 provided support to JM.

## Author contributions

Conceptualization, J.M.B.; Methodology, E.V., J.M., J.M.B.; Formal analysis, E.V., P.B. S.R.M., K.S.; Investigation, E.V., P.B., S.R.M, K.S., A.R.T.; Writing – Original Draft, J.M.B.; Writing – Review & Editing, J.M.B, P.B, E.V., S.R.M., S.U., J.M.; Visualization, E.V., P.B., S.R.M., J.M.B; Funding Acquisition, S.U., J.M.B, A.R.T., J.M; Supervision, S.U., J.M., J.M.B.

## Competing interests

The authors declare no competing interests.

## Materials and methods

### Animal models

Mouse studies were carried out in accordance with UK Home Office regulations and the UK Animals (Scientific Procedures) Act of 1986 (ASPA) under a UK Home Office licence (PPL 70/8719) and approved by the King’s College London Ethical Review Committee. *Nestin-Cre* mice (obtained from Jackson, stock number 003771) and *Tsc1*^*fl/fl*^ mice (obtained from Jackson, stock number 005680) were maintained in a C57BL/6 background. Mice were housed under a 12 hour light/dark cycle with *ad libitum* access to food and water. *Tsc1*^*fl/fl*^ mice were genotyped using primers 5’-GTC ACG ACC GTA GGA GAA GC-3’ and 5’-GAA TCA ACC CCA CAG AGC AT-3’. *Nestin-Cre* mice were genotyped using primers 5’-TTGCTAAAGCGCTACATAGGA-3’ and 5’-GCCTTATTGTGGAAGGACTG-3’.

Pregnant dams were injected intraperitoneally with 1mg/kg rapamycin (Fluorochem Ltd) at E15.5, embryos removed at E16.5, embryonic brains dissected and immediately frozen on dry ice.

### Cell culture, immunoprecipitation and western blot analysis

SH-SY5Y, Neuro-2a (N2a) and HEK293 cells were maintained in Dulbecco’s Modified Eagle’s Medium (DMEM) high glucose (Sigma-Aldrich) supplemented with 10% (v/v) foetal bovine serum (FBS; Sigma-Aldrich) and 1% (w/v) penicillin/streptomycin (Sigma-Aldrich). Cells were trypsinized and sub-cultured twice a week and incubated in 5% CO_2_ at 37°C throughout the experiments unless stated otherwise.

For insulin and rapamycin treatment cells were seeded at 7 × 10^5^ cells per well in a six-well plate (Thermo Scientific) in DMEM + 10% FBS, then serum-starved for 16 hours in DMEM alone. Unless stated otherwise, cells were then incubated in DMEM ±1μM insulin (Sigma-Aldrich), or DMEM + 10% FBS ± 1μM rapamycin (LC Laboratories), or another inhibitor, for one hour. 10μM S6K inhibitor (PF4708671, Sigma-Aldrich), 1μM mTOR kinase active site inhibitor (KU0063794, Sigma-Aldrich) and DL001 (Aeovian Pharmaceuticals Inc. at the concentrations shown in Supplemental Figure 1B) were used. Cells were lysed with 0.2% (w/v) SDS (Sigma-Aldrich), 10mM EDTA, 1x protease inhibitor cocktail (Roche), and 1x phosphatase inhibitor cocktail (Sigma-Aldrich) unless stated otherwise.

C57BL/6J mouse primary cortical cells were extracted at E14.5 and one million cells were plated per well in a six well plate in Neurobasal medium (2% (v/v) B27, 2mM glutamine, 0.3% (w/v) glucose, 37.5mM NaCl, 5% (v/v) FBS; ThermoFisher). 48 hours later the media was replaced with 5% (v/v) FBS ± 1μM rapamycin for one hour and lysed as above.

For lambda protein phosphatase treatment cells were serum starved overnight, stimulated with DMEM ± 1μM insulin, then lysed in 50mM Tris pH 7.5, 150mM NaCl, 0.1% (v/v) Triton, 1X protease inhibitor cocktail ± lambda protein phosphatase (New England Biolabs) as per the manufacturer’s instructions and incubated at 30°C for one hour.

For glucose and amino acid studies, SH-SY5Y cells were seeded and serum starved as above. This was followed by a 2.5 hour incubation in Hanks Balanced Salt Solution (HBSS, Sigma-Aldrich) to starve cells of amino acids and to reduce the concentration of glucose (to 1g/l) and in some conditions a 30 minute pre-incubation of 1μM rapamycin. Trophic factors were then added to the starved cells for 25 minutes with a combination of glucose, amino acids, FBS (all from Sigma-Aldrich), DMEM no glucose (ThermoFisher), insulin (1μM), or rapamycin (1μM), in continued HBSS or DMEM high glucose medium.

For western blot analysis cell extracts were denatured and reduced in 1x sample buffer (500mM Tris pH 6.8, 40% (v/v) glycerol, 0.2% (w/v) SDS, 2% (v/v) β-mercaptoethanol, and 0.02% (w/v) bromophenol blue) and boiled at 98°C for 10 minutes. Proteins were separated by SDS-polyacrylamide gel electrophoresis (PAGE). Proteins were then transferred onto nitrocellulose membranes (GE Healthcare Life Sciences). Membranes were blocked in 10% fat-free milk powder in tris buffered saline (TBS) (50mM Tris pH 7.4, 150mM NaCl) and probed overnight at 4°C with primary antibodies in 5% (w/v) bovine serum albumen (BSA; Fisher) TBS-T (TBS + 0.1% (v/v) Tween 20). Following four 10 minute washes in TBS-T, membranes were incubated with the appropriate horseradish peroxidase-conjugated secondary antibodies (1:4000) in 5% milk TBS-T for one hour at room temperature. Finally, blots were treated with enhanced chemiluminescence reagents (ECL, GE Healthcare) and imaged using a Kodak or Bio-Rad ChemiDoc system imaging system.

Embryonic brains were lysed in urea lysis buffer (8M urea, 50mM HEPES pH8.2, 10mM glycerol-2-phosphate, 50mM NaF, 5mM sodium pyrophosphate, 1mM EDTA, 1mM sodium orthovanadate, 1mM DTT, cOmplete Protease inhibitor cocktail, Sigma phosphatase inhibitor cocktail III, 500nM okadaic acid) and sonicated briefly. Protein concentration was measured using Pierce BCA Assay kit. Twenty micrograms of brain lysate per embryo were denatured and reduced for 10min at 70°C in 1X Pierce LDS sample buffer supplemented with 100mM DTT. Denatured samples were loaded onto pre-cast NuPAGE 4-12%, Bis-Tris protein gel (Invitrogen). Proteins were then transferred onto PVDF membrane using transfer buffer (25mM tris pH8.3, 192mM glycine, 20% v/v methanol) for 1.5hrs at 310mA. Membranes were blocked in 10% w/v skimmed-milk (Marvel) dissolved in TBS-T for 1 hour.

Membranes used for immunoblotting with phospho-specific antibodies were washed with TBS-T prior to incubation at 4°C overnight with primary antibodies in 5% w/v BSA dissolved in TBS-T. Overnight membranes were then washed three times with TBS-T and incubated with an appropriate HRP-conjugated secondary antibody diluted in 5% w/v milk. After washing the PVDF membranes with TBS-T, they were incubated with enhanced ECL solution (Cytiva) and imaged using Amersham Imager 600 (GE Healthcare). Western blot images were quantified using the Amersham Imager 600 Software.

Primary antibodies used were rabbit anti-Unkempt (1:1000, HPA023636, Cambridge Bioscience), rabbit anti-mTOR (1:1000, #2972, Cell Signalling), rabbit anti-Raptor (1:1000, #2280, Cell Signalling), rabbit anti-β-actin (1:5000, #4967, Cell Signalling), rabbit anti-S6K (1:1000, #9202, Cell Signalling), rabbit anti-phosphoS6K (Thr389, 1:1000, #9234, Cell Signalling), rabbit anti-S6 (1:1000, #2217, Cell Signalling), rabbit anti-phospho-S6 (Ser235/236, 1:1000, #2211, Cell Signalling), rabbit anti-4E-BP1 (1:1000, #9644, Cell Signalling), rabbit anti-phospho-4E-BP1 (Thr37/46, 1:1000, #2855, Cell Signalling), rabbit anti-FLAG (1:2000, F7425, Sigma-Aldrich), rat anti-FLAG (1:2000, #200473-21, Agilent), mouse anti-V5 (1:1000, R960-25, ThermoFisher), mouse anti-HA (1:1000, sc-7392, Santa Cruz), rabbit anti-phosphoUnkempt (1:200, S606, S611, this study), rabbit anti-phospho AKT (Ser473, 1:1000, 4060T, Cell Signalling), rabbit anti-AKT (1:1000, 5691T, Cell Signalling), mouse anti-α-tubulin (1:10000, T9026, clone DM1A, Sigma-Aldrich), rabbit anti-hamartin (TSC1) (1:1000, ab270967, clone EPR24364-109, Abcam), rabbit anti-PKCα (1:1000, ab32376, clone Y124, Abcam), rabbit anti-phospho-PKCα (Ser657, 1:1000, ab180848, clone EPR1901(2), Abcam).

### Expression vectors, cloning, derivation of inducible cell lines, and transfections

Full length mouse Unkempt was amplified from cDNA using primers 5’-CACCAGATATCCAATGTCGAAGGGCCCCGGGCCCG-3’ and 5’-GACGACTCTAGATCACGACTGGAGGGCATGGGCCC-3’ and cloned into pENTR/D-TOPO (ThermoFisher) according to the manufacturer’s instructions to create pENTR-Unk. Primers used to generate the Unkempt deletions and zinc finger constructs in pENTR-Unk were the same as used previously (Murn *et al*., 2015). Primers used to generate the SerL deletion in pENTR-Unk were 5’-GGGCCAGGGGCTGCTGAGCT-3’ and 5’-GCCCACAGGGAGGATGCCCA-3’ and to generate the SerS deletion in pENTR-Unk were 5’-GGGCCAGGGGCTGCTGAGCT-3’ and 5’-GTTATCAAAAGTCTTCTCTAGG-3’. These constructs were then recombined into pcDNA3.1/nV5-DEST (ThermoFisher). pCAG-Rheb^CA^ (also known as pCAG-Rheb^S16H^), was a gift from Angelique Bordey (Hartman *et al*., 2013).

The alanine and aspartate substitution mutations in Unkempt were created by PCR, using the SuperFi mastermix (Thermo Fisher) as per the manufacturer’s instructions using pENTR-Unk as a template. The PCR products were digested using DpnI (NEB) at 37°C for 90 minutes and then transformed into DH5α chemically competent cells. The following primers were used:

S467-S479>A (S467A, S474A, S475A, S476A, T478A, S479A)

Forward primer – 5’

GACTTCCAGCATCGCTGCCGCTATTGCCGCCAGCTTGGCAGCCACTCCC 3’

Reverse primer – 5’

GCTGGAGGTAATAGCGGCAGCGATGCTGGAAGTCAGGGGGGCGCCCACAGG 3’

S467-S479>D (S467D, S474D, S475D, S476D, T478D, S479D)

Forward primer – 5’

GACTTCCAGCATCGATGACGATATTGACGACAGCTTGGCAGCCACTCCC 3’

Reverse primer – 5’

GCTGTCGTCAATATCGTCATCGATGCGGAAGTCAGGGGGTCGCCCACAGG 3’

T491A

Forward primer – 5’ CTGCAGGCGCCAACAGCACCCCTGGCATGAATGC 3’

Reverse primer – 5’ GCTGTTGGCGCCTGCAGGGCTGGGGGGAG 3’

T491D

Forward primer – 5’ CTGCAGGCGACAACAGCACCCCTGGCATGAATGC 3’

Reverse primer – 5’ GCTGTTGTCGCCTGCAGGGCTGGGGGGAG 3’

S546A

Forward primer – 5’ CACCCCAGCGCCGTCACAATCGGTGGC 3’

Reverse primer – 5’ CGATTGTGACGGCGCTGGGGTGGGGCACTGC 3’

S546D

Forward primer – 5’ CACCCCAGCGACGTCACAATCGGTGGC 3’

Reverse primer – 5’ CGATTGTGACGTCGCTGGGGTGGGGCACTGC 3’

S565-S578>A (S565A, S571A, S574A, S576A, S578A)

Forward primer – 5’

CAGCTCAGCTGCCTTCCACGCTGCTGCTCCAGCCCCTCCCGTCAGCCTCTCC 3’

Reverse primer -5’

GGGAGGGGCTGGAGCAGCAGCGTGGAAGGCAGCTGAGCTGCCCAGGGCGCCAG GGATG 3’

S565-S578>D (S565D, S571D, S574D, S576D, S578D)

Forward primer – 5’

CAGCTCAGCTGACTTCCACGATGCTGATCCAGACCCTCCCGTCAGCCTCTCC 3’

Reverse primer – 5’

GGGAGGGTCTGGATCAGCATCGTGGAAGTCAGCTGAGCTGCCCAGGTCGCCAGG GAT 3’

S596-S611>A (S596A, S598A, T605A, S606A, S608A, S611A)

Forward primer – 5’

CGTTTTTGGGGGCCGCAGCAGCACATGGAGCTTTGGGTCTGAACGGG 3’

Reverse primer - 5’

CCAAAGCTCCATGTGCTGCTGCGGCCCCCAAAAACGTGTTTTCTGCCTGAGCCAA GTGGCC 3’

S596-S611>D (S596D, S598D, T605D, S606D, S608D, S611D)

Forward primer – 5’

CGTTTTTGGGGGACGACGCAGACCATGGAGATTTGGGTCTGAACGGG 3’

Reverse primer – 5’

CCAAATCTCCATGGTCTGCGTCGTCCCCCAAAAACGTGTTTTCGTCCTGATCCAA GT 3’

S631-S640>A (S631A, T634A, S635A, S640A)

Forward primer – 5’

CCAGGCGCTGCCCCTGCCTTCCTAGCAGGGCCAGGGGCTGC 3’

Reverse primer -5’

CCCTGCTAGGAAGGCAGGGGCAGCGCCTGGGGCGAAGCTTCCAGAGGC 3’

S631-S640>D (S631D, T634D, S635D, S640D)

Forward primer – 5’

CCAGGCGATGACCCTGCCTTCCTAGACGGGCCAGGGGCTGC 3’

Reverse primer – 5’

CCCGTCTAGGAAGGCAGGGTCATCGCCTGGGTCGAAGCTTCCAGAGGC 3’

Doxycycline-inducible HeLa cells used for the analyses of cellular morphogenesis and for CLIP experiments were generated as described previously (Murn *et al*., 2015). Briefly, stably rtTA3-expressing cells were infected ecotropically with retroviruses for TREtight-driven expression of GFP alone (pTt-IGPP) or GFP and either wild-type Unkempt protein or any of the Unkempt alanine or aspartate substitution mutants indicated in Figure 6 and created as described above (pTtight-X-IGPP, where X is either wild-type or mutant Unkempt). To induce transgene expression, double-selected cells were treated with doxycycline (Sigma-Aldrich) at 1 ug/ml.

For transient transfections, HEK293 cells were seeded at 2 ×10^5^ cells per well in six well plates (Thermo Fisher) in DMEM/10% FBS. The following day, cells were transfected with V5-tagged Unkempt plasmids at 1 μg/well, using polyethylenimine (Sigma) at 7 μg/well and 200 μl/well of Optimem (Sigma). For co-transfections, Rheb^CA^ was used at 0.5 μg/well and the V5-tagged Unkempt constructs at 1 μg/well. Cells were lysed 48 hours post transfection using 0.2% (w/v) SDS (Sigma-Aldrich), 10mM EDTA, 1x protease inhibitor cocktail (Roche), and 1x phosphatase inhibitor cocktail (Sigma-Aldrich).

### Co-immunoprecipitation experiments and liquid chromatography with tandem mass spectrometry (LC-MS/MS) analysis

To induce overexpression of and immunoprecipitate FLAG-HA-Unkempt, previously described HeLa S3 cells stably expressing doxycycline-inducible FLAG-HA-Unkempt (Murn *et al*., 2015) were used. They were cultured in DMEM (high glucose; Sigma) supplemented with 10% (v/v) FBS (Sigma) and 1% penicillin/streptomycin (Lonza) in 15 cm dishes (Nunc, ThermoFisher) to 90% confluency and induced with doxycycline hyclate (Sigma-Aldrich) at 1 μg/ml for 24 hours. Non-induced cells were also cultured in parallel to use as a negative control. 16 hours prior to co-immunoprecipitation, cells were serum starved in DMEM, in the presence of doxycycline. 10% FBS was then re-introduced to culture media and cells were treated with 1 μM rapamycin (LC Laboratories) or DMSO for one hour. Dishes were then washed with ice-cold PBS and cells were scraped and lysed on ice for 15 minutes in 3 ml lysis buffer.

For co-immunoprecipitations, one 15 cm dish was used for each condition and cells were lysed in either CHAPS lysis buffer (120 mM NaCl, 40 mM HEPES pH 7.5, 1 mM EDTA, 0.3% (w/v) CHAPS, 1X protease inhibitor cocktail, 1X phosphatase inhibitor cocktail and 0.2 mM PMSF) or Triton X-100 lysis buffer (120 mM NaCl, 40 mM HEPES pH 7.5, 1 mM EDTA, 1% Triton X-100, 1X protease inhibitor cocktail, 1X phosphatase inhibitor cocktail and 0.2 mM PMSF). Samples were then centrifuged at 17,000 x g for 10 minutes at 4°C. The supernatant was transferred to a new tube and protein concentration was determined using Protein Assay Dye Reagent Concentrate (Bio-Rad) following the manufacturer’s instructions. Samples were diluted and standardized to the condition with lowest protein concentration (≤ 2 mg/ml) using their respective lysis buffer. Equal volumes of diluted lysates were added to pre-washed 50 μl anti-FLAG M2 affinity agarose gel slurry (Sigma-Aldrich) and incubated rotating end over end for two hours at 4°C. The agarose beads were then washed three times with lysis buffer followed by three additional washes with wash buffer (150 mM NaCl, 50 mM HEPES pH 7.5). Excess buffer was removed using an insulin needle (30G) and beads were resuspended in 60 μl 2 x sample buffer and protein was eluted by heating at 95°C for 5 min. Co-immunoprecipitants were resolved on an 8% SDS-PAGE gel followed by protein transfer on nitrocellulose membrane (GE Healthcare Life Sciences). Western blot analysis was performed as described above.

For LC-MS/MS analysis of immunoprecipitated Unkempt, cells were cultured in two 15 cm dishes for each condition and samples were processed in a similar way to co-immunoprecipitation experiments but using a different lysis buffer (25 mM Tris pH 8.0, 150 mM NaCl, 5% (v/v) glycerol, 1% Triton (v/v) X-100, 1X protease inhibitor cocktail, 1X phosphatase inhibitor cocktail and 0.2 mM PMSF) and wash buffer (50 mM Tris pH 8.0, 150 mM NaCl). Sample dilution and standardization prior to incubation with anti-FLAG resin was omitted. Bound Unkempt was eluted with 40 μl 2x sample buffer heated at 95°C for 5 minutes. 35μl of each eluate was resolved on a 4-12% Bis-Tris Novex NuPAGE gel (ThermoFisher) for coomassie staining and mass spec analysis. The remaining 5 μl were resolved on the same gel and used for FLAG western blotting to confirm the immunoprecipitation and exact molecular weight of Unkempt. Half of the gel for LC-MS/MS analysis was fixed (7% (v/v) glacial acetic acid, 40% (v/v) methanol, 53% HPLC-grade water) for 30 minute shaking at room temperature. The fixed gel was stained overnight at room temperature (4 parts 1X Brilliant Blue-G Colloidal Concentrate (Sigma-Aldrich) and 1-part methanol, freshly mixed). The gel was destained three to four times (5% (v/v) glacial acetic acid, 25% (v/v) methanol, 70% HPLC-grade water) until background was significantly reduced and bands were clearly visible. Bands of the expected FLAG-HA-Unkempt size (∼90 kDa) for both conditions, rapamycin and DMSO, were excised using sterile scalpel in a fume hood and stored in HPLC-grade water.

LC-MS/MS analysis was performed by the Cambridge Centre for Proteomics, University of Cambridge. Gel fragments were transferred into a 96-well PCR plate. The bands were cut into 1mm^2^ pieces, destained with a solution 50% acetonitrile and 50mM ammonium bicarbonate, reduced with 10mM DTT for one hour at 37°C and alkylated with 55mM iodoacetamide at room temperature in the dark for 45 minutes, then subjected to enzymatic digestion with 0.01μg/μl sequencing grade chymotrypsin (Promega V1061) overnight at 37°C. After digestion, the supernatant was pipetted into a sample vial and loaded onto an autosampler for automated LC-MS/MS analysis.

All LC-MS/MS experiments were performed using a Dionex Ultimate 3000 RSLC nanoUPLC (Thermo Fisher Scientific Inc, Waltham, MA, USA) system and a QExactive Orbitrap mass spectrometer (Thermo Fisher Scientific Inc, Waltham, MA, USA). Separation of peptides was performed by reverse-phase chromatography at a flow rate of 300 nL/minute and a Thermo Scientific reverse-phase nano Easy-spray column (Thermo Scientific PepMap C18, 2 μm particle size, 100A pore size, 75 μm i.d. x 50cm length). Peptides were loaded onto a pre-column (Thermo Scientific PepMap 100 C18, 5 μm particle size, 100A pore size, 300 μm i.d. x 5mm length) from the Ultimate 3000 autosampler with 0.1% (v/v) formic acid for three minutes at a flow rate of 10 μL/minute. After this period, the column valve was switched to allow elution of peptides from the pre-column onto the analytical column. Solvent A was water + 0.1% (v/v) formic acid and solvent B was 80% (v/v) acetonitrile, 20% water + 0.1% (v/v) formic acid. The linear gradient employed was 2-40% B in 30 minutes.

The LC eluant was sprayed into the mass spectrometer by means of an Easy-Spray source (Thermo Fisher Scientific Inc.). All *m/z* values of eluting ions were measured in an Orbitrap mass analyzer, set at a resolution of 70000 and was scanned between *m/z* 380-1500. Data dependent scans (Top 20) were employed to automatically isolate and generate fragment ions by higher energy collisional dissociation (HCD, NCE:25%) in the HCD collision cell and measurement of the resulting fragment ions was performed in the Orbitrap analyser, set at a resolution of 17500. Singly charged ions and ions with unassigned charge states were excluded from being selected for MS/MS and a dynamic exclusion window of 20 seconds was employed.

Post-run, the data was processed using Protein Discoverer (version 2.1., ThermoFisher). Briefly, all MS/MS data were converted to mgf files and the files were then submitted to the Mascot search algorithm (Matrix Science, London UK) and searched against the UniProt_Mus_musculus_20180514 database (61295 sequences; 27622875 residues). The peptide and fragment mass tolerances were set to 20 ppm and 0.1 Da, respectively. A significance threshold value of p<0.05 and a peptide cut-off score of 20 were also applied. Following Mascot analysis, data were converted into .sf3 file and sent to user for further analysis.

Converted files were analysed using Scaffold4 Proteome Software (version 4.8.7). Protein threshold was set to 95% with minimum number of peptides at 2. Peptide threshold was relaxed and each phospho-peptide was examined manually accounting for accurate identification of amino acids, number of peptides with the same phosphorylation site and peptide identification probability percentage. All selected sites were less than 10 delta ppm.

Prediction disorder probability was performed using the online tool Protein DisOrder prediction System (PrDOS, http://prdos.hgc.jp/cgi-bin/top.cgi) (Ishida & Kinoshita, 2007) applying 5% prediction false positive rate. Mouse Unkempt protein sequence was used. Protein sequences were aligned using Clustal Omega.

### In vitro mTORC1-Rheb kinase assays

HeLa S3 cells stably expressing doxycycline inducible FLAG-HA-Unkempt were used (Murn *et al*., 2015). After serum starvation, stimulation and rapamycin treatment, as described above, cells from three 15 cm dishes per condition were scraped and pooled together. They were lysed for 20 min on ice in 10 ml lysis buffer (25 mM Tris pH 8.0, 150 mM NaCl, 5% (v/v) glycerol, 1% (v/v) Triton X-100, 1X protease inhibitor cocktail, 1X phosphatase inhibitor cocktail and 0.2 mM PMSF). Lysates were centrifuged at 20,000 x g for 20 minutes at 4°C and the cleared supernatant was incubated with 70 μl anti-FLAG M2 affinity agarose gel rotating end over end overnight at 4°C. Samples were then washed twice in lysis buffer followed by three washes in high salt buffer (50 mM Tris pH 8.0, 500 mM NaCl, 5% (v/v) glycerol, 1% (v/v) Triton X-100) and three times in wash buffer (50 mM Tris pH 8.0, 150 mM NaCl). Recombinant FLAG-HA-Unkempt was eluted in 80 μl wash buffer supplemented with 0.2 mg/ml 3 x FLAG peptide (Generon) rotating end over end for 30 min at 4°C. Purified Unkempt was collected using 30G needle, flash-frozen in liquid nitrogen and stored at -80°C until required.

Radiolabelled *in vitro* mTORC1 kinase assays with Rheb-GTP were performed as described previously (Dunlop *et al*., 2009) using FLAG-HA-Unkempt purified as described above.

For the *in vitro* kinase assays for LC-MS/MS analysis, HEK293 cells grown in twelve 10 cm dishes were co-transfected with active myc-mTOR E2914K mutant (Addgene) and HA-Raptor (Kim *et al*., 2002), each at 9 μg per plate. In parallel another eight 10 cm dishes of HEK293 cells were transfected with 18 μg of the constitutively active mutant GST-FLAG-Rheb Q64L (Rheb^CA^) (Long *et al*, 2005). 48 hours later cells co-transfected with mTOR and Raptor were serum starved overnight and then stimulated with 100 nM insulin (Sigma-Aldrich) for 15 minutes. Cells were washed in ice-cold PBS and each plate was lysed in 1 ml mTOR lysis buffer (40 mM HEPES pH 7.4, 50 mM NaCl, 2 mM EDTA, 10 mM B-glycerophosphate, 0.3% (w/v) CHAPS) on ice for 10 minutes. Samples were centrifuged for 10 minutes at 17,000 x g at 4°C. Cleared supernatants were pooled together and incubated with 3 μl anti-c-Myc antibody (9E10, DSHB) per 1 ml lysate, rotating end over end for 1.5 hours at 4°C. The lysate was divided equally and incubated with 50 μl of prewashed 50% Protein G slurry (GE Healthcare) for each condition, rotating for one hour at 4°C. Beads were then washed once with low salt mTOR wash buffer (40mM HEPES pH 7.4, 150mM NaCl, 2mM EDTA, 10mM B-glycerophosphate, 0.3% (w/v) CHAPS), twice with high salt mTOR wash buffer (40mM HEPES pH 7.4, 400mM NaCl, 2mM EDTA, 10mM B-glycerophosphate, 0.3% (w/v) CHAPS), and twice with HEPES/KCl wash buffer (25mM HEPES pH 7.4, 20mM KCl). Cells transfected with GST-FLAG-Rheb Q64L were washed in ice-cold PBS and each plate was lysed in 1 ml Rheb lysis buffer (40 mM HEPES pH 7.4, 50 mM NaCl, 10 mM pyrophosphate, 10 mM glycerophosphate, 5 mM MgCl_2_, 0.3% (w/v) CHAPS, 1X protease inhibitor cocktail) for 10 minutes on ice followed by centrifugation for 10 minutes at 17,000 xg at 4°C. Supernatants were incubated with 30 μl prewashed Glutathione sepharose 4B slurry (GE Healthcare) per 1 ml lysate, rotating end over end for two hours at 4°C. Beads were then washed twice with Rheb lysis buffer and once with Rheb storage buffer (20 mM HEPES pH 8.0, 200 mM NaCl, and 5 mM MgCl_2_). The supernatant was removed with 30G needle and bound GST-Rheb was eluted in 40 μl Rheb elution buffer (20 mM HEPES pH 8.0, 200 mM NaCl, and 5 mM MgCl_2_, 10 mM reduced Glutathione). To prepare the mTORC1 complex inhibitor, 5 μg FKBP12 was incubated in 50 μl of 30 mM rapamycin in 1x kinase assay buffer (25mM HEPES pH 7.4, 20mM KCl, 10mM MgCl_2_) for five minutes in the dark at room temperature. For each condition, mTOR/Raptor immunoprecipitates (bound on beads) were incubated with 50 μl 3x kinase assay buffer, 25 μl GST-FLAG-Rheb Q64L, with or without 10 μl FKBP12:rapamycin complex, made up to 150 μl total reaction volume and incubated on ice for 20 minutes. The *in vitro* kinase assay was started by adding 50 μl assay start buffer (25mM HEPES pH 7.4, 10mM MgCl_2_, 140mM KCl, 500uM ATP) and 1 μg purified FLAG-HA-Unkempt, or recombinant GST-4E-BP1 (Dunlop *et al*., 2009) as a positive control. Samples were incubated for one hour shaking at 30°C. The reaction was stopped with 4 x sample loading buffer and stored at -20°C. For mass spectrometry analysis, kinase assay reaction samples were diluted in 1 x kinase assay buffer/assay start buffer (3:1) and concentrated using Amicon Ultra-0.5 centrifugal filter unit with a 3 kDa cutoff. Concentrated samples were processed and analysed by LC-MS/MS as described above. Positive control samples were used for western blotting to confirm mTORC1 activity as assessed by phosphorylation of 4EBP-1. Primary antibodies used were mouse anti-c-Myc (1:1000, 9E10, DSHB), mouse anti-HA (1:1000, SC-7392, Santa Cruz), rat anti-FLAG (1:2000, #200473-21, Agilent), rabbit anti-4E-BP1 (1:1000, #9644, Cell Signalling), rabbit anti-phospho-4E-BP1 (Thr37/46, 1:1000, #2855, Cell Signalling).

### UV-crosslinking and immunoprecipitation (CLIP) assays

CLIP experiments were carried out essentially as described in (Buchbender *et al*, 2020). GFP or GFP and Unkempt-inducible HeLa cells were grown in 10 cm plates and harvested at 80% confluency. Prior to harvest, the inducible HeLa cells were treated with doxycycline for 24 hours to induce transgene expression. Cells were washed, irradiated with UV-light (254 nm) at 150 mJ/cm^2^ on ice, scraped into three 2 ml tubes, and spun down at 4°C. The supernatant was removed and the cell pellets were frozen at -80°C until use. Immunoprecipitation of the cross-linked Unkempt-RNA complexes was carried out using anti-Flag antibody (F1804, Sigma-Aldrich). Samples were processed exactly as described in (Buchbender *et al*., 2020) until the autoradiography stage where the membrane was exposed to X-ray film (the 3’ end dephosphorylation and first adapter ligation steps were omitted). The experiment was performed in three independent biological replicates. All replicates yielded comparable results.

### Unkempt phospho-specific antibody generation

Three phospho-peptides were generated corresponding to residues 601-610 (containing phospho-serine 606), 607-615 (containing phospho-serine 611) and 601-615 (phospho-serines 606 and 611) in Unkempt. Rabbits were immunised with all three phospho-peptides and phospho-specific antibodies were affinity purified using the phospho-peptide corresponding to residues 601-615 (containing phospho-serines 606 and 611). Antibody generation was performed by Covalab.

### Morphogenesis assays

Cellular morphologies were analysed as described previously (Schreiber *et al*., 2019). Briefly, the morphology of each cell was evaluated by an axial ratio (y/x) where the length of the absolute longest cellular axis (y) was divided by the length of the longest axis perpendicular to the y-axis (x). Lengths of x-and y-axes of GFP-positive inducible HeLa cells were measured at 48 hours of treatment with doxycycline. At least 50 GFP-positive cells were quantified for each inducible cell line.

### Statistical analyses

Data were analysed using GraphPad Prism version 7 (GraphPad Software). Data distributions were assessed for normality, then analysed using a Student’s t-test. p values < 0.05 were considered significant.

### Materials and Data Availability

All methods and data are described in the manuscript or supplemental materials.

## Notes

### Competing Interest Statement

The authors have declared no competing interest.

