## Supplemental figures for "Phosphorylation of the novel mTOR substrate Unkempt regulates cellular morphogenesis"

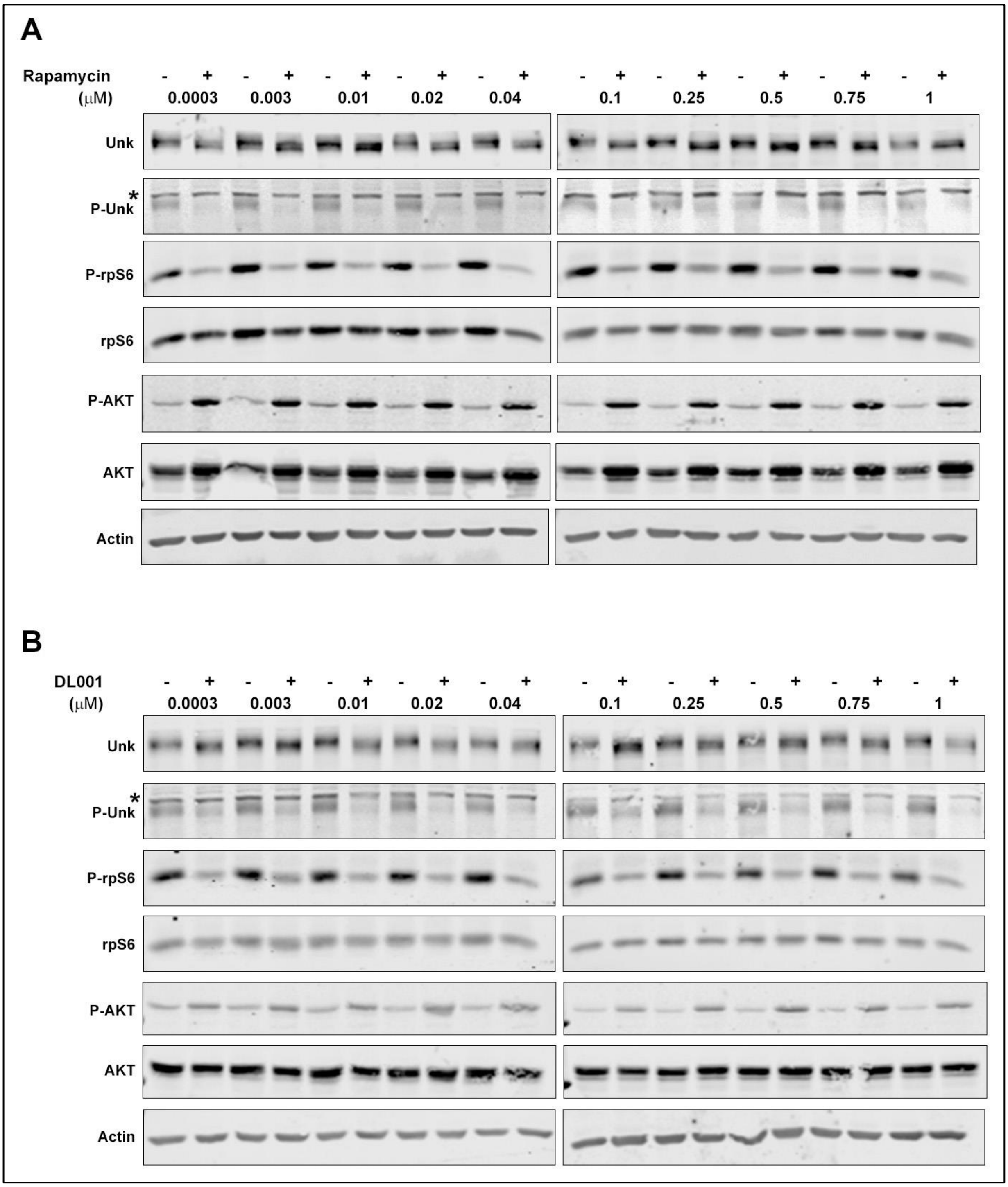


*Figure S1. Unkempt phosphorylation is highly sensitive to mTORC1 inhibition.* (A) The electrophoretic mobility of Unkempt in SH-SY5Y cells is decreased by a wide range of rapamycin concentrations. P-Unk is probed using an Unkempt phospho-S606/phospho-S611 specific antibody, see text for details. *indicates a non-specific band. (B) Unkempt phosphorylation in SH-SY5Y cells is inhibited by a wide of concentration of the highly selective mTORC1 inhibitor DL001. *indicates a non-specific band.

```
Mm      MSKGPGPGGSAASSAPPAATAQVLQAQPEKPQHYTYLKEFRTEQCPLFVQHKCTQHRPYT    60
Hs      MSKGPGPGGSAASSAPPAATAQVLQAQPEKPQHYTYLKEFRTEQCPLFVQHKCTQHRPYT    60
Dm      -------------MLANETNKLLLSSQQEKPNHYTYLKEFRVEQCQSFLQHKCNQHRPFV    47
                       .  :.  :*.:* ***:*********.***  *:****.****:.

Mm      CFHWHFVNQRRRRSIRRRDGTFNYSPDVYCTKYDEATGLCPEGDECPFLHRTTGDTERRY    120
Hs      CFHWHFVNQRRRRSIRRRDGTFNYSPDVYCTKYDEATGLCPEGDECPFLHRTTGDTERRY    120
Dm      CFNWHFQNQRRRRPVRKRDGTFNYSADNYCTKYDETTGICPEGDECPYLHRTAGDTERRY    107
        **:*** ******.:*:********.* *******:**:********:****:*******

Mm      HLRYYKTGICIHETDSKGNCTKNGLHCAFAHGPHDLRSPVYDIRELQAMEALQNGQTTVE    180
Hs      HLRYYKTGICIHETDSKGNCTKNGLHCAFAHGPHDLRSPVYDIRELQAMEALQNGQTTVE    180
Dm      HLRYYKTCMCVHDTDSRGYCVKNGLHCAFAHGMQDQRPPVYDIKEL---ETLQNAESTLD    164
        ******* :*:*:***:* *.*********** :* *.*****:**   *:***.::*::

Mm      GSIEGQSAGAASHAMIEKILSEEPRWQETAYVLGNYKTEPCKKPPRLCRQGYACPYYHNS    240
Hs      GSIEGQSAGAASHAMIEKILSEEPRWQETAYVLGNYKTEPCKKPPRLCRQGYACPYYHNS    240
Dm      STNALNALDKE-----RNLMNEDPKWQDTNYVLANYKTEPCKRPPRLCRQGYACPQYHNS    219
        .:   :: .       .:::.*:*:**:* ***.********:************ ****
                      #
Mm      KDRRRSPRKHKYRSSPCPNVKHGDEWGDPGKCENGDACQYCHTRTEQQFHPEIYKSTKCN    300
Hs      KDRRRSPRKHKYRSSPCPNVKHGDEWGDPGKCENGDACQYCHTRTEQQFHPEIYKSTKCN    300
Dm      KDKRRSPRKYKYRSTPCPNVKHGEEWGEPGNCEAGDNCQYCHTRTEQQFHPEIYKSTKCN    279
        **:******:****:********:***:**:** ** ***********************
                                           #                       #
Mm      DMQQAGSCPRGPFCAFAHIEPPPLSDDVQPSSAVSSPTQPGPVLYMPSAAGDSVPVSPSS    360
Hs      DMQQSGSCPRGPFCAFAHVEQPPLSDDLQPSSAVSSPTQPGPVLYMPSAAGDSVPVSPSS    360
Dm      DVQQAGYCPRSVFCAFAHVEPCSMDD----------------------------------    305
        *:**:* ***. ******:*  .:.*

Mm      PHAPDLSALLCRNSGLGSPSHLCSSPPGPSRKASNLEGLVFPGESSLAPGSYKKAPGFER    420
Hs      PHAPDLSALLCRNSSLGSPSNLCGSPPGSIRKPPNLEGIVFPGESGLAPGSYKKAPGFER    420
Dm      -----------------------------------------PRENSLS------------    312
                                                 * *..*:

Mm      EDQVGAEYLKNFKCQAKLKPHSLEPRSQEQPLLQPKQDVLGILPVGSPLTSSISSSITSS    480
Hs      EDQVGAEYLKNFKCQAKLKPHSLEPRSQEQPLLQPKQDMLGILPAGSPLTSSISSSITSS    480
Dm      ---------------ASLANTSLLTRSS-APINIPN----------TTLSNSINDFNSGS    346
                       *.*   ** .**.  *:  *:          :.*:.**..  :.*

Mm      LAATPPSPAGTNSTPGMNANALPFYPTSDTVESVIESALDDLDLNEFGVAALEKTFDNSA    540
Hs      LAATPPSPVGTSSVPGMNANALPFYPTSDTVESVIESALDDLDLNEFGVAALEKTFDNST    540
Dm      FAVNIPS------------SSLTYSPTN--------------HANLFNVDAFNYGGSN-K    379
        :*.. **            .:*.: **.              . * *.* *::   .*

Mm      VPHPSSVTIGGSLLQSSAPVNIPGSLGSSASFHSASPSPPVSLSSHFLQQPQGHLSQSEN    600
Hs      VPHPGSITIGGSLLQSSAPVNIPGSLGSSASFHSASPSPPVSLSSHFLQQPQGHLSQSEN    600
Dm      LSNSLSATQNDSSLFFPSRIISPG-FG-----DGLSISPSVRIS---------ELNTIRD    424
        :.:. * * ..* *  .: :  ** :*     .. * **.* :*         .*.  .:
               #  #
Mm      TFLGTSASHGSLGLNGMNSSIWEHFASGSFSPGTSPAFLSGPGAAELARLRQELDEANGT    660
Hs      TFLGTSASHGSLGLNGMNSSIWEHFASGSFSPGTSPAFLSGPGAAELARLRQELDEANST    660
Dm      DINSSSVGN-SLFENTLNT------AKNAFSLQS----LQSQNNSDLGRITNELLTKNAQ    473
         : .:*..: **  * :*:      *..:**  :    *.. . ::*.*: :**   *.

Mm      IKQWEESWKQAKQACDAWKKEAEEAGERASAAGAECELAREQRDALELRVKKLQEELERL    720
Hs      IKQWEESWKQAKQACDAWKKEAEEAGERASAAGAECELAREQRDALEVQVKKLQEELERL    720
Dm      IHKLNG---------------------RFEDMACKLKIAELHRDKAKQEAQEWKERYD--    510
        *:: :                      * .  ..: ::*. :**  : ..:: :*. :

Mm      HTVPEAQTLPAAPDLEALSLSTLYSIQKQLRVHLEQVDKAVFHMQSVKCLKCQEQTR--A    778
Hs      HAGPEPQALPAFSDLEALSLSTLYSLQKQLRAHLEQVDKAVFHMQSVKCLKCQEQKR--A    778
Dm      ---LAQIQLNLPAELRDLSIQKLKQLQSKLRTDLEEVDKVLYLENAKKCMKCEENNRTVT    567
                *   .:*. **:..* .:*.:**..**:***.::  :: **:**:*:.*  :

Mm      VLPCQHAVLCELCAE-GSECPVCQPSRAHALQS    810
Hs      VLPCQHAALCELCAE-GSECPICQPGRAHTLQS    810
Dm      LEPCNHLSICNTCAESVTECPYCQVPVITTHT-    599
        : **:*  :*: ***  :*** **     :
```

*Figure S2. Phosphorylated residues identified in Unkempt.* The primary sequence of *Mus musculus* (Mm) Unkempt aligned with *Homo sapiens* (Hs) and *Drosophila melanogaster* (Dm) Unkempt. Grey highlighted residues were phosphorylated in vehicle control and rapamycin treatment conditions; blue highlighted residues were phosphorylated only in the vehicle control condition; green highlighted residues were phosphorylated only in the rapamycin treatment condition. # indicates residues phosphorylated only in the control condition in the mTORC1 *in vitro* kinase assay. See Supplemental Figure S3 for details.

| | IP HeLa cells | | | |
|---|---|---|---|---|
| | No. times identified | | *In vitro* kinase assay | |
| Residue | Control | Rapamycin | Control | Rapamycin/ FKBP12 |
| S85 | 2 | 1 | | |
| S255 | 2 | 2 | YES | NO |
| S332 | 0 | 2 | | |
| S335 | 3 | 3 | YES | YES |
| S336 | 2 | 2 | YES | NO |
| T338 | 3 | 3 | YES | YES |
| S357 | 3 | 3 | YES | YES |
| S359 | 2 | 3 | YES | NO |
| S360 | 3 | 3 | | |
| S367 | 2 | 1 | | |
| S378 | 3 | 3 | YES | YES |
| S380 | 0 | 1 | | |
| S385 | 2 | 2 | | |
| S467 | 1 | 1 | | |
| S474 | 2 | 1 | | |
| S475 | 0 | 1 | | |
| S476 | 0 | 1 | | |
| T478 | 1 | 0 | | |
| S479 | 2 | 2 | | |
| T491 | 1 | 0 | | |
| S546 | 0 | 1 | | |
| S565 | 1 | 0 | | |
| S571 | 1 | 1 | | |
| S574 | 1 | 0 | | |
| S576 | 1 | 3 | | |
| S578 | 3 | 3 | YES | YES |
| S596 | 1 | 0 | | |
| S598 | 3 | 3 | | |
| T605 | 0 | 1 | YES | YES |
| S606 | 2 | 2 | | |
| S608 | 1 | 0 | YES | NO |
| S611 | 3 | 0 | YES | NO |
| S631 | 3 | 1 | | |
| T634 | 0 | 1 | | |
| S635 | 1 | 0 | | |
| S640 | 1 | 0 | | |

*Figure S3. mTORC1 dependent and independent phosphorylated residues in Unkempt*. Columns 1-3 show a summary of the number of times each phosphorylated residue was identified by LC-MS/MS in FLAG-HA-Unkempt immunoprecipitated from vehicle (DMSO) control and rapamycin treated HeLa S3 cells in three biological replicates. Columns 4 and 5 show phosphorylated residues identified by LC-MS/MS of purified FLAG-HA-Unkempt following an *in vitro* kinase assay with reconstituted mTORC1/Rheb$^{CA}$ in control or rapamycin/FKBP12 conditions.

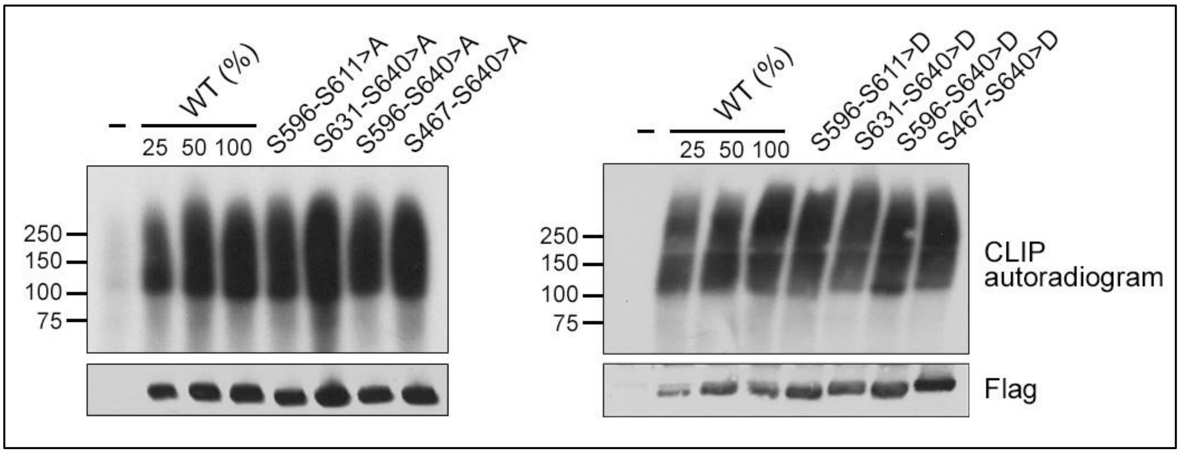


*Figure S4. Phosphosite mutants of Unkempt retain gross RNA-binding capacity.* Binding of the indicated Flag-HA-tagged alanine (>A) (left) or aspartate (>D) (right) mutants of Unkempt or wild-type (WT) Unkempt to RNA in inducible HeLa cells was analyzed by CLIP at 24 hours of treatment with or without (-) doxycycline. Top, CLIP autoradiograms; bottom, immunoblots of the membrane used to develop the autoradiograms. Total alanine and aspartate mutant and (-) samples (100%) were used for the analysis
